# Distinct classes of antidepressants commonly act to shape pallidal structure and function in mice

**DOI:** 10.1101/2024.09.23.614626

**Authors:** Yoshifumi Abe, Yuki Sugiura, Rae Maeda, Shu Taira, Keisuke Yoshida, Daisuke Ibi, Satoru Moritoh, Kenji Hashimoto, Sho Yagishita, Kenji F Tanaka

## Abstract

Antidepressants including selective serotonin reuptake inhibitors, ketamine, and psilocybin are all effective for treating depression despite their distinct primary mechanisms. We hypothesized that these drugs may share a common mechanism that underlies their therapeutic actions. We treated mice with one of the following: escitalopram, *R-*/*S*-/*RS-*ketamine, or psilocin. Additionally, groups exposed to electroconvulsive stimulation and a saline control were included. Following treatment, fixed brains underwent structural magnetic resonance imaging, and voxel-based morphometry was performed to evaluate brain-wide volumetric changes. Compared with control treatment, we observed greater volumes in the nucleus accumbens, ventral pallidum, and external globus pallidus across all antidepressant treatments, and a smaller volume in the mediodorsal thalamus. Specifically, *R*-ketamine, *RS*-ketamine, and psilocin induced more pronounced hypertrophy of the ventral pallidum, whereas selective serotonin reuptake inhibitors and *S*-ketamine predominantly increased the volume of the external globus pallidus. Further analyses using super-resolution microscopy and imaging mass spectrometry revealed corresponding microstructural and molecular changes. Greater pallidal volume was associated with striatal medium spiny neuron terminal hypertrophy and elevated γ-aminobutyric acid (GABA) levels. Interestingly, all antidepressants were also associated with higher striatal dopamine content. Moreover, striatal vesicular GABA transporter overexpression reproduced the medium spiny neuron terminal hypertrophy and increased pallidal GABA content, and was associated with a reduction in innate anxiety. These findings indicate that despite their pharmacological diversity, antidepressant treatments lead to shared pallidum-centered structural and molecular changes. We propose that these shared changes may potentiate the striato-pallidal inhibitory circuit, thereby contributing to the overall antidepressant effect.

## Introduction

Antidepressant treatments are a substantial therapeutic option for major depressive disorder, and selective serotonin reuptake inhibitors (SSRIs) are used as a first-line treatment ^1^. SSRIs immediately block serotonin (5-HT) transporters and enhance 5-HT signaling, but long-term SSRI use is necessary ^2^. By contrast with first-line antidepressants, the dissociative anesthetic ketamine (*N*-methyl-D-aspartate receptor antagonist) ^3, 4^ and the psychedelic psilocybin (5-HT_2A_ receptor agonist) ^5^ have recently been recognized as rapid-acting antidepressants offering long-term effects from a single dose. Although the primary pharmacological targets and temporal modes of action differ among these types of antidepressants, they can similarly induce recovery states in patients with major depressive disorder, indicating common underlying mechanisms for the induction and maintenance of recovery. This indication is also supported by previous studies that aimed to address the hidden shared mechanisms across distinct medications ^6–10^.

The acquisition of a recovery state can be understood as an outcome of plastic changes in the brain from a diseased state to a healthy state after treatment. Such brain state changes likely involve unidentified cellular changes, functional plastic changes in the circuitry, and structural plastic changes. Because structure always correlates with function, we hypothesize that identifiable structural plasticity should accompany theoretical functional plasticity. Our strategy involved screening for brain regions undergoing structural plastic changes using structural magnetic resonance imaging (MRI), followed by closer inspection of those regions with super-resolution microscopy (SRM) to identify specific structural plastic changes.

Using this strategy, we studied naïve mice with well-validated innate anxiety-like behaviors that can be alleviated with SSRI, ketamine, or psilocin (the active metabolite of psilocybin) treatment. We revealed that all antidepressants were associated with increases in pallidum volume, hypertrophy of vesicular γ-aminobutyric acid (GABA) transporter (VGAT)-positive terminals of striatal medium spiny neurons (MSNs), and increased GABA content in these terminals. Striatal VGAT overexpression reproduced the MSN terminal hypertrophy and increased pallidal GABA content, and was associated with a reduction in innate anxiety. Additional volumetric and histological analysis indicated that antidepressants can be divided into two subtypes. We found that the ventral pallidum (VP) and mediodorsal thalamus (MD) were hubs for volumetric changes. Therefore, the nucleus accumbens (NAc)–VP–MD axis was highlighted as an antidepressant action-related circuitry.

## Methods and Materials

### Animals

All animal procedures were conducted in accordance with the National Institutes of Health Guide for the Care and Use of Laboratory Animals and approved by the Animal Research Committee of Keio University School of Medicine (A2022-029). Experiments were performed using 8- to 20-week-old male C57BL/6J mice. All mice were maintained on a 12-h/12-h light/dark cycle (lights on at 08:00), and behavioral experiments were conducted during the light phase. C57BL/6J mice (8 weeks old) were purchased from Oriental Yeast Co., Ltd. (Tokyo, Japan).

### Antidepressant treatment

Psilocin was synthesized within the Japanese law according to the methods published by Shirota et al.^11^, and its purity was verified by nuclear magnetic resonance analysis (Avance III 600, Bruker, MA, USA). R-ketamine and S-ketamine were prepared by recrystallization of RS-ketamine (Ketalar®, ketamine hydrochloride, Daiichi Sankyo Pharmaceutical Ltd., Tokyo, Japan), as described previously^12^. Escitalopram was used in the SSRI experiments and was purchased from Tokyo Chemical Industry Co., Ltd. (Tokyo, Japan). Psilocin, ketamine, and escitalopram were dissolved in saline.

Mice received a single acute intraperitoneal (i.p.) injection of vehicle, psilocin, SR-ketamine, R-ketamine, or S-ketamine. On the basis of previous studies^13–15^, the dose was 1 mg/kg for psilocin and 10 mg/kg for R-ketamine or S-ketamine. RS-ketamine was mixed using 5 mg/kg of R-ketamine and 5 mg/kg of S-ketamine. The mice were perfused 1 week after a single injection with vehicle, psilocin, or ketamine. Escitalopram (15 mg/kg per day in deionized water) or vehicle was delivered by oral gavage for 3 weeks ^13^. The mice were perfused 1 day after the final administration of escitalopram.

### ECS

The procedure was previously described ^16^. Briefly, ECS was administered to mice that were anesthetized with 3.0% sevoflurane once daily for 3 weeks (three times per week, for a total of nine ECS sessions). ECS was administered in square wave pulses of 25 mA and 0.5 msec pulse width for 1 second at a frequency of 100 Hz via bilateral ear clip electrodes (UgoBasile, Comerio, Italy). The control mice underwent the same procedure on the same schedule without electrical stimulation. The mice were perfused 1 day after the final ECT administration.

### Plasmid construction and adeno-associated virus (AAV) preparation

AAV vectors were prepared and purified as previously reported ^17^. The transfer plasmids (pAAV-hSyn-ALFA-VGAT-pA or pAAV-CaMKII (0.3)-EGFP-miR30-shRNA VGAT-WPRE-pA)^6^ and packaging plasmids (pHelper (Stratagene) and pAAV2/5 which was a gift from Melina Fan (Addgene plasmid # 104964)) were co-transfected into Lenti-X 293T cells (Takara Bio INC., Shiga, Japan) using polyethyleneimine (23966-100; Polysciences).

After 7 days, AAVs were collected from the cell lysate and supernatant and purified by ultracentrifugation using iodixanol. The purified solution was concentrated by ultrafiltration using an Amicon Ultra-15 (NMWL 30 K; Merck Millipore, MA, USA). Virus titer (GC/mL) was measured by quantitative PCR (primer set AAV2ITR-F/AAV2ITR-R or WPRE-F/WPRE-R) using THUNDERBIRD Next SYBR qPCR Mix (TOYOBO Co., Ltd., Osaka, Japan) on a StepOne real-time PCR system (Applied Biosystems, MA, USA). The following AAV vectors were used in the experiments: AAV_5_-hSyn-ALFA-VGAT-pA (6.9×10^13^ GC/mL), AAV_5_-CaMKII-EGFP-miR30-shRNA VGAT-WPRE-pA (3.3×10^13^ GC/mL), and AAV5-hSyn-EGFP-pA (#50465, Addgene).

### Viral vector injection

AAVs were injected into wild-type mice. Mice were anesthetized with a mixture of ketamine and xylazine (i.p., 100 mg/kg and 10 mg/kg, respectively) and fixed in a stereotaxic apparatus (Narishige, Tokyo, Japan). The skull surface was exposed, and the periosteum and blood were removed. Next, 0.3 μL of viral solution was injected into the right dorsal striatum (+0.6 mm anteroposterior and ±2.2 mm mediolateral from bregma and −2.2 mm dorsoventral from the brain surface according to the atlas of Paxinos and Franklin) and NAc (+1.0 mm anteroposterior and ±1.4 mm mediolateral from bregma and −4.0 mm dorsoventral from the brain surface) through a small hole using a glass micropipette connected to a Nanoliter 2020 injector (World Precision Instruments, FL, USA). The mice were used 4 weeks after the AAV injection.

### Corticosterone (CORT)-induced depression model mice

First, 70 μg/mL CORT (Tokyo Chemical Industry Co., Ltd., Tokyo, Japan) was dissolved in 0.45% b-cyclodextrin (Sigma, St. Louis, MO, USA) in water ^18^. CORT (10 mg/kg/day) was delivered in the drinking water for 6 weeks, and SSRI (escitalopram) treatment started after 3 weeks of CORT treatment. Escitalopram (15 mg/kg per day) or vehicle was delivered by oral gavage for 3 weeks.

### Behavioral tests

We performed the open field (OF) test on day 1, the novelty suppressed feeding (NSF) test on day 2, and the elevated plus maze (EPM) test on day 3. For wild-type mice treated with antidepressants, the behavioral tests were performed either 1 week after the single injection of ketamine or psilocin or the day after the last SSRI treatment (Fig, 1L). For mice with VGAT manipulation, the behavioral tests were performed 4 weeks after AAV injection (Fig. 5A).

In the OF test (day 1), mice were tested in the OF (a 50 × 50 cm square; Melquest Ltd., Toyama, Japan) for 30 min. The center region was defined as the inner 25 × 25 cm area. The light intensity in the field was set to 30 lux. The behavior of each mouse was recorded from above using a video camera. The percentage time spent in the center area was calculated using in-house software written with MATLAB (MathWorks, Natick, MA, USA). After testing, the mouse was returned to its home cage and food deprived for 1 day before the NSF test.

The NSF test (day 2) was performed in the same field as the OF test. Light intensity in the field was set to 1000 lux. A single food pellet (the same food as in the home cage) was placed on green paper (an 8 × 8 cm square) in the center of the OF. Each mouse was placed in a corner of the OF, and the latency to begin eating the food pellet was recorded (up to a maximum of 10 min).

In the EPM test (day 3), mice were tested in an EPM apparatus (Melquest) for 20 min. The maze had arms that were 6 cm wide, 56 cm long, and 43 cm above the floor. Two opposing closed arms had walls that were 15 cm high. Light intensity in the open arms only was set to 800 lux. The behavior of each mouse was recorded from above using a video camera. The percentage time spent in the open arm was calculated using in-house software written with MATLAB.

### MRI

The procedures used in the *ex vivo* MRI study were previously described ^6, 16^. Mice were deeply anesthetized with a mixture of 0.3 mg/kg medetomidine, 4 mg/kg midazolam, and 5 mg/kg butorphanol and perfused with a 4% paraformaldehyde phosphate buffer solution. The brains were removed along with the skull and postfixed in the same fixative for 24 h. The fixed brains were then stored in PBS for 1 week. The duration of paraformaldehyde and PBS immersion was consistent for all samples to avoid differences in brain volumes caused by postperfusion immersion fixation and storage ^19^. Brains within their skulls were firmly fixed using fitted sponges into an acrylic tube (22-mm diameter) filled with Fluorinert (Sumitomo 3M Limited, Tokyo, Japan) to minimize the signal intensity attributable to the embedding medium. Additionally, vacuum degassing was performed to reduce air bubble-derived artifacts in the structural images. The MRI study was performed with an 11.7 T BioSpec 117/11 US/R unit (Biospin GmbH, Ettlingen, Germany) and a volume-type coil with a 23-mm inner diameter for transmitting and receiving. For brain volumetric analysis, the structural images were acquired using 3D-T2-weighted multi-slice rapid acquisition with relaxation enhancement (RARE) with the following parameters: repetition time=2000 ms, echo time=75 ms, spatial resolution=100×100×100 µm, RARE factor=32, and averages=8. The structural images after treatment with saline, psilocin, ketamine, and SSRIs were obtained in the current study, and those of brains with ECT treatment and controls were obtained from a previous study using the same protocol^16^ and were reanalyzed in this study.

To check image quality, we calculated the signal-to-noise ratio of the T2 images (Supplementary Fig. 1A). The signal-to-noise ratio was quantified by dividing the mean signal intensity within a homogeneous white matter (WM) region of the brain by the standard deviation of the background noise ^20^.

### Voxel-based morphometry (VBM) analysis

Image preprocessing and statistical analysis for the whole-brain voxel-based analysis were performed using SPM12 (Wellcome Trust Centre for Neuroimaging, London, UK) and in-house software written using MATLAB (MathWorks, Natick, MA, USA). First, each T2-weighted image was resized by a factor of 10 to account for the whole-brain volume differences between humans and rodents and aligned to the same space by registering each image to tissue probability maps (TPMs). Using the Allen Mouse Brain Atlas as a guide, we generated the TPMs with T2 data from a separate cohort of 12 eight[week[old male mice. Each T2 image was segmented into gray matter (GM), WM, and cerebrospinal fluid (CSF) using a unified segmentation approach, which enabled bias correction, image registration to the TPMs, and tissue classification. The segmented GM, WM, and CSF images were then merged to create a brain mask. Using this brain mask, skull stripping was performed on the T2 images, and the masked images were segmented again into GM, WM, and CSF (Supplementary Fig. 1B). We carefully checked that landmark regions—including WM areas (e.g., the corpus callosum and anterior commissure) and CSF compartments (e.g., the lateral ventricles)—were appropriately delineated. All subjects in the control, psilocin-, ketamine-, SSRI-, and ECS-treated groups were then pooled, and a study[specific GM template was generated using the Diffeomorphic Anatomical Registration Through Exponentiated Lie Algebra algorithm. The second sets of segmented images were then spatially normalized into this study[specific GM template, and modulated GM images were obtained for each animal by applying the determinant of the Jacobian of the transformation, thereby accounting for the expansion and/or contraction of brain regions. These images were smoothed with a 3-mm (corresponding to 0.3 mm in the original resolution) full width at half maximum Gaussian kernel. The total brain volume was defined as a sum of the GM volume (GMV) and the WM volume.

Whole-brain voxel-wise comparisons of the GMV between each treated mouse and its control were performed using SPM12 with the total brain volume as a covariate. The first analysis compared the GMV between antidepressant- or ECS-treated mice and control mice. The significance threshold for the voxel-wise whole-brain analysis was set at a cluster-level family-wise error corrected p value < 0.05, with an individual voxel threshold of an uncorrected p value = 0.001. The significant clusters were overlayed on the averaged T2 image.

Clustering of volumetric changes was performed by principal component analysis. Statistical t-values were calculated from comparisons of the brain volume values of each defined subregion (see below) between antidepressant-treated mice and control mice. The t-values across the defined subregions were used for principal component analysis.

### ROI-based structural covariance analysis

To define ROIs of brain subregions, we utilized the Turone Mouse Brain Template and Atlas (TMBTA) ^21^, which is available at https://www.nitrc.org/projects/tmbta_2019/, and modified the brain subregions. A total of 397 brain subregions were defined in each hemisphere. The individual masked T2 images for all subjects were non-linearly registered to the TMBTA T2 image using the *antsRegistrationSyN* script of Advanced Normalization Tools (ANTs) (available at https://github.com/ANTsX/ANTs). The inverse transforms from the TMBTA T2 image of each subject were then applied to register the TMBTA atlas to each subject (Supplementary Fig. 1C). Next, the brain volumes of each defined subregion were estimated by averaging the voxel values (the determinant value of the Jacobian transformation) of each region.

Structural covariance analysis was performed using the NBS toolbox (https://www.nitrc.org/projects/nbs/) by reference to previous studies ^22–24^. Structural covariance was quantified using the partial correlation coefficient between paired brain volume value estimates across each antidepressant group. This approach yielded a separate connectivity matrix with dimensions of 794×794 for each group that quantified the connectivity strength between all pairs of regions (Fig. 5A). These structural covariances were subjected to Fisher transformation to obtain the z-transformed values. The delta value was calculated by dividing the difference in z-values by the standard deviation (SD) of the difference in z-values. This calculation was performed independently for each pair of regions. The SD of the difference in z-values was estimated by bootstrapping the sample 1000 times. Edges in the network matrices that were 2.5 or more times the SD were identified as suprathreshold. The threshold of SD ≥ 2.5 is equal to p < 0.05 because the difference in z-values follows a normal distribution. The identified regions were overlayed on the mouse template brain (Fig. 5B). To evaluate the characteristics of the extracted network, we estimated the degree of centrality and betweenness centrality of the nodes. Nodes with centralities greater than 1 SD from the mean were defined as hubs of the network.

### Deformation-based morphometry (DBM) analysis

The procedures of this analysis were referred to the previous study ^25^. Each skull-stripped T2 image was corrected for intensity non-uniformity using the *N3BiasFieldCorrection* script of ANTs. A study-specific T2 template was then constructed from the corrected T2 images of all subjects in the control, psilocin-, ketamine-, SSRI-, and ECS-treated groups using the *antsMultivariateTemplateConstruction2* script of ANTs. Jacobian determinant maps were calculated from the flow fields in the T2 template using the *CreateJacobianDeterminantImage* script of ANTs, and were log-scaled to allow the voxel-wise estimation of apparent volume change. To compare volume differences between antidepressant- or ECS-treated mice and appropriate control mice, voxel-based statistics were performed on the log-transformed Jacobian determinant maps using SPM12, with the total brain volume as a covariate. The significance threshold for the voxel-wise whole-brain analysis was set at a cluster-level family-wise error-corrected p-value < 0.05, with an individual voxel threshold of an uncorrected p-value = 0.001. Significant clusters were overlaid on the study-specific T2 template.

### Histology

After the MRI study, brains were used for histological assessments. The brains were cryoprotected in 20% sucrose overnight and frozen. Frozen brains were cut on a cryostat at 40-µm thickness for the floating sections. The sections were incubated with primary antibodies overnight at room temperature. The following antibodies were used: anti-VGAT (1:1000 dilution; guinea pig polyclonal, Frontier Institute, Hokkaido, Japan, Cat# MSFR106160), anti-Bassoon (1:1000 dilution; mouse monoclonal, SAP7F407, Enzo Life Sciences, Farmingdale, NY, USA, Cat# ADI-VAM-PS003-F), anti-Gephyrin (1:1000 dilution, mouse monoclonal, mAb7a, Synaptic Systems, Coventry, UK, Cat# 147021), anti-GFP (1:250 dilution, goat polyclonal; Rockland Immunochemicals, Pottstown, PA, USA, Cat# 600-101-215), anti-ALFA (1:500 dilution; rabbit polyclonal, NanoTag, Göttingen, Germany, Cat# N1581), anti-NeuN (1:1000 dilution, rabbit monoclonal, EPR12763, Abcam, Cambridge, UK, Cat# ab190565), anti-VGluT1 (1:1000 dilution; rabbit polyclonal, Frontier Institute, Cat# MSFR106190), and anti-VGluT2 (1:1000 dilution; rabbit polyclonal, Frontier Institute, Cat# MSFR106310). The sections were then treated with species-specific secondary antibodies conjugated to Alexa Fluor 488, 555, or 647 for 2 h at room temperature. Mounting medium (ProLong Glass Antifade Mountain, Thermo Fisher Scientific, Waltham, MA, USA) was applied to the samples, and they were mounted with coverslips (thickness: No. 1, Matsunami Glass). Macro-fluorescence images were obtained using an inverted microscope (BZ-X710; Keyence, Osaka, Japan). Micro-fluorescence images were obtained using super-resolution microscopy (SRM).

### Super-resolution microscopy

Structured illumination microscopy (SIM)-SRM micro images were obtained using a Zeiss ELYRA 3D-SIM system equipped with an EM-CCD camera (Carl Zeiss). Before obtaining the SIM-SRM images, precise alignment for the different wavelengths of light was performed using the same mounting medium (ProLong Glass) containing 0.1% Tetraspeck (0.2-µm beads, Thermo Fisher Scientific) to correct for unavoidable laser misalignment and optical aberrations, which can cause the alignment not to coincide at high resolution. Next, 14–20 z-section images were obtained at intervals of 130 nm using a 64× objective lens. The number of pattern rotations of the structured illumination was adjusted to three in the ELYRA system. After obtaining all images, the SIM images were reconstructed and aligned using the channel alignment data.

### Analysis of SRM images

This analysis was performed in a blind manner. SRM images were obtained as previously described^6^ and analyzed using ImageJ software (http://rsb.info.nih.gov/ij/). The optimal brightness and grayscale pixel values were manually determined to provide the sharpest discrimination of the microstructure border. These adjusted images were then converted into binary images. To calculate the density of VGAT^+^, VGLuT1^+^, VGluT2^+^, Bassoon^+^, or Gephyrin^+^ puncta, SRM images of each type of staining were randomly obtained at three locations each within the GPe, VP, MD, and LHb. The numbers of puncta were counted, and the mean number over the three randomly selected locations was calculated. To calculate puncta size, 3D-SRM images of each type of staining were randomly obtained at three locations in each region. We observed the 3D image and used the single plane at which the size was largest to calculate the size. We measured the area of 150 puncta from each of the three locations. The mean size of the 150 puncta was used as the representative puncta size for each animal.

To calculate the size of the NeuN^+^ soma of GPe and VP neurons, 3D-SRM images of NeuN staining were randomly obtained within the GPe and VP. Although NeuN is predominantly a nuclear marker, the soma can also be visualized with NeuN staining ^26^. We observed the 3D image and used the single plane at which the size was largest to calculate the size. We calculated the size of 20–30 somas per mouse and used the average size as the representative soma size for each animal.

### Mass spectrometry imaging

The procedures were previously described^6^. Mice were deeply anesthetized with a mixture of 0.3 mg/kg medetomidine, 4 mg/kg midazolam, and 5 mg/kg butorphanol, and the brains were rapidly removed from the skull and frozen in liquid N_2_. Next, 10-μm-thick sections of fresh-frozen brains were prepared with a cryostat and thaw-mounted on conductive indium-tin-oxide-coated glass slides (Matsunami Glass). A pyrylium-based derivatization method was applied for the tissue localization imaging of neurotransmitters^27, 28^. TMPy (4.8 mg; Taiyo Nippon Sanso Co., Tokyo, Japan) was dissolved in a mixed solution (methanol:water:TEA=70:25:5) and was applied to brain sections using an airbrush (Procon Boy FWA Platinum 0.2-mm caliber airbrush, Mr. Hobby, Tokyo, Japan). To enhance the reaction efficiency of TMPy, TMPy-treated sections were placed into a dedicated container and allowed to react at 60°C for 10 min. The container included two channels in the central partition to wick moisture from the wet filter paper region to the sample section region. The filter paper was soaked with 1 mL of methanol/water (70%/30% volume/volume) and placed next to the section inside the container, which was then completely sealed to maintain humidity levels. The TMPy-labeled brain sections were sprayed with matrix (2,5-dihydroxybenzoic acid in 70% ethanol) using an automated pneumatic sprayer (TM-Sprayer, HTX Tech., Chapel Hill, NC, USA). Ten passes were performed according to the following conditions: flow rate, 120 µL/min; airflow, 10 psi; nozzle speed, 1100 mm/min.

To detect the laser spot area, the sections were scanned, and laser spot areas (200 shots) were detected with a spot-to-spot center distance of 80 µm. Signals between *m*/*z* 100–650 were corrected. The section surface was irradiated with yttrium aluminum garnet laser shots in the positive ion detection mode using matrix-assisted laser desorption/ionization time-of-flight mass spectrometry (MALDI-TOF MS; timsTOF fleX, Bruker Daltonics, Bremen, Germany). The laser power was optimized to minimize the in-source decay of targets. The mass spectrometry spectra obtained were reconstructed to produce mass spectrometry images using Scils Lab software (Bruker Daltonics). Optical images of brain sections were obtained using a scanner (GT-X830, Epson, Tokyo, Japan), followed by MALDI-TOF MS of the sections. The detected mass of TMPy-labeled standard GABA (*m*/z 208.163) was increased by 105.0 Da compared with the original mass (molecular weight 103.0 Da). Tandem mass spectrometry was used to confirm the fragmentation ions of TMPy from the standard sample. A fragmented ion of the pyridine ring moiety (*m/z* 122.1) was regularly cleaved and observed for all TMPy-modified target molecules.

### Statistical analysis

Statistical processing was performed using MATLAB. Two-tailed Student’s t-tests were performed to compare treated and control mice. Bonferroni corrections were applied to correct p-values for multiple comparisons. Values are shown as the mean and standard error of the mean and are plotted as scatter diagrams.

## Results

### The pallidum is a common region for antidepressant-mediated volume increases

To identify common structural plasticity changes after antidepressant treatment, we treated mice with antidepressants at concentrations and times that have previously demonstrated antidepressant effects, including the SSRI escitalopram (daily administration for 3 weeks, 15 mg/kg) ^13^, *R*-ketamine, *S*-ketamine, racemic (*SR*) ketamine (single injection, 10 mg/kg) ^13^, or psilocin (single injection, 1 mg/kg) ^14^ (Fig. 1A). We then conducted structural MRI and an unbiased, brain-wide volumetric analysis known as VBM 1 week after the single injection of ketamine and psilocin or 3 weeks after the initial SSRI treatment.

**Fig. 1:**
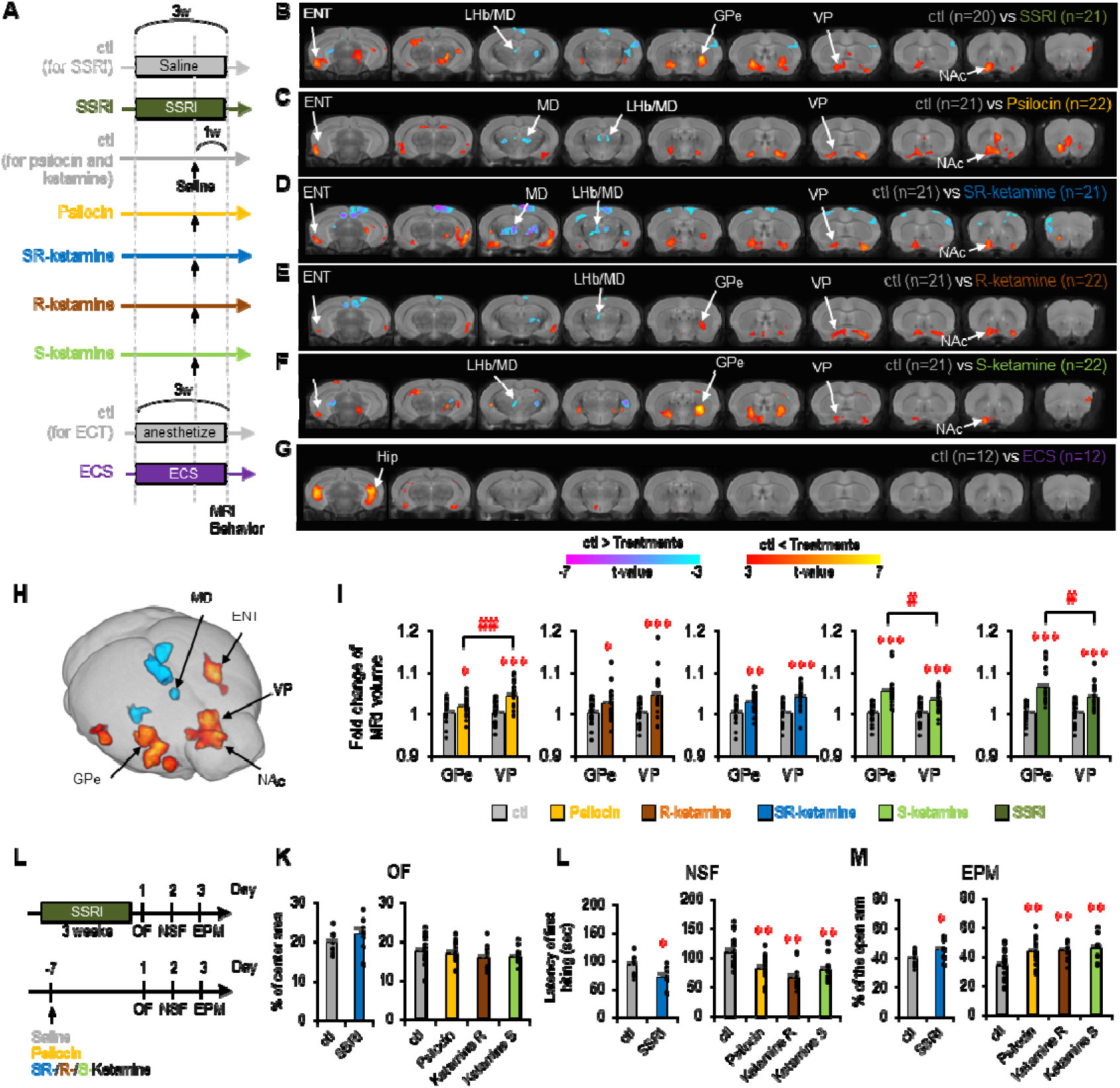
The pallidum is a common region for antidepressant-mediated volume increases in mice. (A) Experimental time course for the administration of saline, psilocin, ketamine, escitalopram (SSRI), or electroconvulsive stimulation (ECS). (B–G) Voxel-based morphometry (VBM) analysis of brain volume differences between treated mice and the appropriate control mice (ctl). Hot colors indicate significant volume increases compared with those in the ctls (family-wise error corrected *p* < 0.05), and cool colors indicate the opposite. (H) Volume changes in overlapping regions of the brain after treatment. Red indicates commonly increased volumes, and blue indicates commonly decreased volumes. (I) Fold changes in the brain volume of the globus pallidus (GPe) and ventral pallidum (VP) after each antidepressant treatment. \**p* < 0.05, \*\**p* < 0.01, \*\*\**p* < 0.001 (Student *t* tests were used for comparisons with ctls, and *p* values are adjusted with post hoc Bonferroni corrections for multiple comparisons). #*p* < 0.05, ##*p* < 0.01 (Student *t* tests were used for comparisons between the GPe and VP of the treated mice, *p* values are presented with post hoc Bonferroni corrections). (J) Experimental time course of behavioral tests. (K–M) Behavioral outcomes of the open field (OF), novelty suppressed feeding (NSF), and elevated plus maze (EPM) tests in the ctl (n = 13), SSRI (n = 9), psilocin (n = 13), *R*-ketamine (n = 10), and ketamine (n = 10) groups. All groups were compared with appropriate control mice. ENT, entorhinal cortex; Hip, hippocampus; LHA, lateral hypothalamus area; LHb, lateral habenula; MD, mediodorsal nucleus of the thalamus; NAc, nucleus accumbens.

After 3 weeks of repeated SSRI treatment, volumes of the ventral striatum, including the nucleus accumbens (NAc), and striatal projection regions, such as the external globus pallidus (GPe), VP, substantia nigra pars reticulata (SNr), and entorhinal cortex (ENT), were increased. Conversely, volumes of the lateral habenula (LHb) and the mediodorsal thalamus (MD) were decreased (Fig. 1B). Surprisingly, a single injection of ketamine or psilocin induced similar volumetric changes in these regions (Fig. 1C–F). Across all antidepressants, common volumetric changes included increases in NAc, VP, GPe, and ENT volumes, as well as decreases in MD volume (Fig. 1H and Supplementary Fig. 2). ROI-based volumetric analyses confirmed these results (Supplementary Fig. 3). Plotting the volume values for the GPe and VP further supported significant volume increases with all antidepressant treatments (Fig. 1I). Additionally, we validated our VBM results by applying an alternative volumetric analysis: DBM analysis ^29–31^. Although there were differences in cluster size between the two methods, DBM provided consistent results (Supplementary Fig. 4A–F), confirming the common volumetric changes in the NAc, VP, and MD across antidepressant treatments (Supplementary Fig. 4G).

Next, to examine whether these antidepressant-associated volumetric changes were distinctive, we compared them with the volumetric changes observed after electroconvulsive stimulation (ECS), an animal model of electroconvulsive therapy (ECT) ^16, 32^. After 9 sessions of ECS over 3 weeks, a volume increase was observed mainly within the hippocampus (Fig. 1G and Supplementary Fig. 4F), suggesting that the volumetric changes induced by antidepressants and ECS are distinct. Taken together, these results indicate that the pallidum volume increases are a common effect of various antidepressants. We focused on the pallidum for further histological analysis.

We then examined whether our antidepressant treatments decreased innate anxiety in wild-type mice by performing the OF, NSF, and EPM tests on mice treated with SSRIs, ketamine, or psilocin following the same regimen as in the MRI studies (Fig. 1L). In the NSF test, the latency to the first bite was significantly lower in all antidepressant-treated mice (Fig. 1L), and the percentage of time spent in the open arms of the EPM was higher (Fig. 1M). By contrast, the percentage of time spent in the center area of the OF test was similar among all groups (Fig. 1K). These results indicate that antidepressant treatment affects innate anxiety.

### Hypertrophy of VGAT-positive MSN terminals and increased GABA levels are common microstructural and molecular changes in antidepressant-treated mice

We next investigated the molecular and anatomical basis for the increase in pallidal volume. We previously demonstrated that the VGAT expression level in dorsal striatal MSNs determines the size of their axon terminals (presynaptic structure) and the size of connecting neuron soma and dendrites (postsynaptic structure) ^6^. Additionally, the convergence of such microstructural changes led to MRI-detectable changes in GPe volume. Considering the analogy between ventral and dorsal striatal projections, we examined the inhibitory pre- and postsynaptic structures in both the VP and GPe after antidepressant treatments using SRM (Fig. 2A, B). All classes of antidepressants induced increases in the size and density of VGAT^+^ puncta (presynaptic, Fig. 2C, D and Supplementary Fig. 5A, B), suggesting that VGAT was overexpressed in MSNs. They also induced an increase in NeuN^+^ soma size in the VP and GPe principal neurons (Fig. 2E and Supplementary Fig. 5C). Additionally, we observed some structural changes in the excitatory presynapses in the GPe and VP (Supplementary Fig. 6). However, these alterations are unlikely to involve macroscopic volume changes due to the low punctal density.

**Fig. 2:**
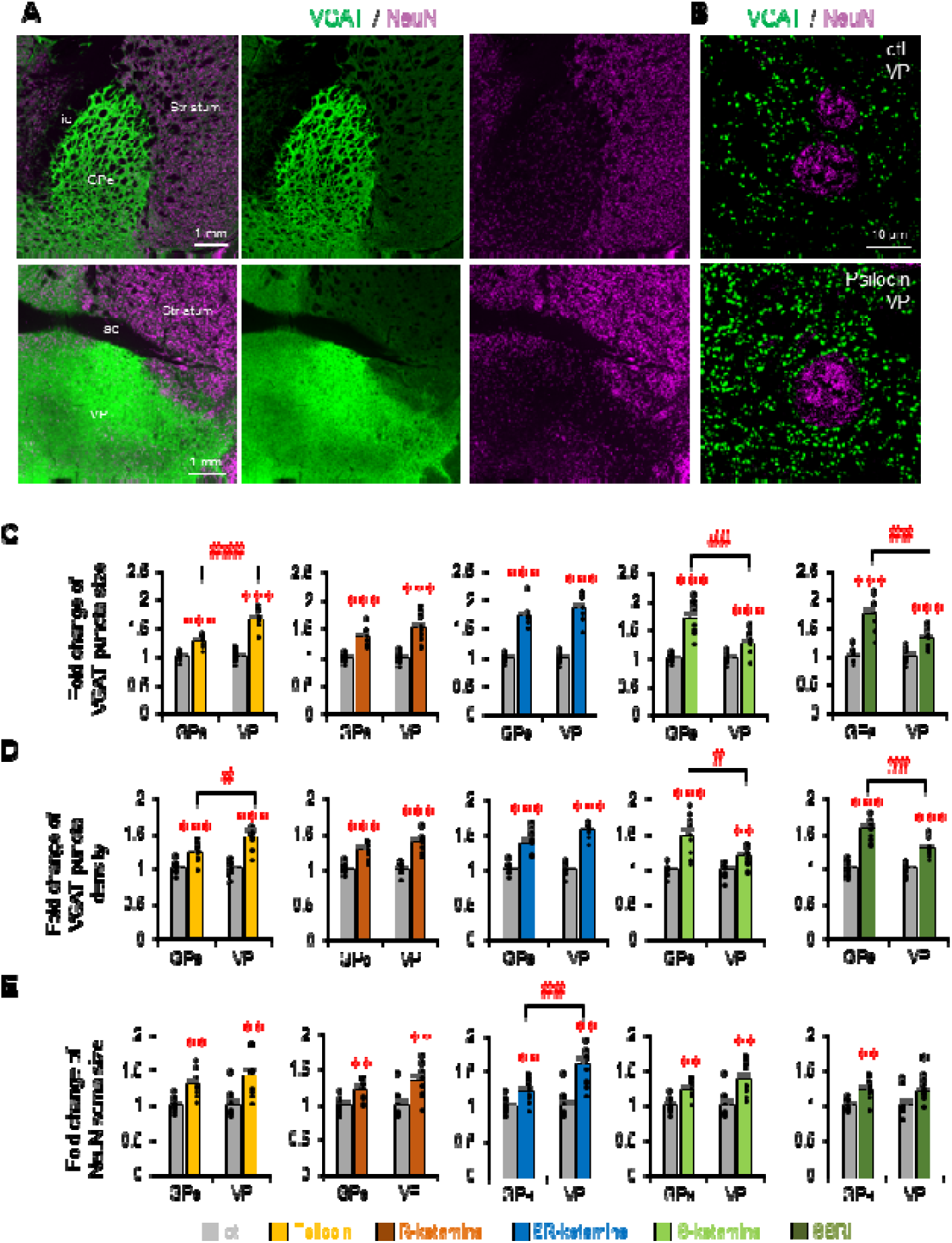
Hypertrophy of MSN terminals and pallidal neuron somas is a common microstructural change in antidepressant-treated mice. (A) Representative microscopy images including the globus pallidus (GPe) and the ventral pallidum (VP). The VGAT and NeuN are labeled in green and magenta, respectively. ic, internal capsule; ac, anterior commissure. (B) Representative super-resolution microscopy images of VGAT and NeuN staining in the VP of control (ctl) and psilocin-treated mice. (C–E) Fold changes in the size and density of VGAT puncta in medium spiny neuron (MSN) terminals and in the size of GPe/VP neuronal soma after antidepressant treatment (n = 10 mice for each treatment). \**p* < 0.05, \*\**p* < 0.01, \*\*\**p* < 0.001 (Student *t* tests were used for comparisons with each ctl, and *p* values are presented with post hoc Bonferroni corrections for multiple comparisons). #*p* < 0.05, ##*p* < 0.01, ###*p* < 0.001 (Student *t* tests were used for comparisons between the GPe and VP of the treated mice, *p* values are presented with post hoc Bonferroni correction). The values are plotted as the means ± standard errors of the means.

Our previous research showed that MSN terminal hypertrophy is coupled with increased GABA levels ^6^. Therefore, we hypothesized that antidepressants might similarly increase GABA levels in the basal ganglia. Using imaging mass spectrometry (IMS), we quantified GABA levels. As expected, treatment with all classes of antidepressants resulted in increased GABA content in the striatum, GPe, and VP (Fig. 3A, B). These results indicate that all antidepressants commonly induce VGAT overexpression, MSN terminal hypertrophy, and increased GABA levels.

**Fig. 3:**
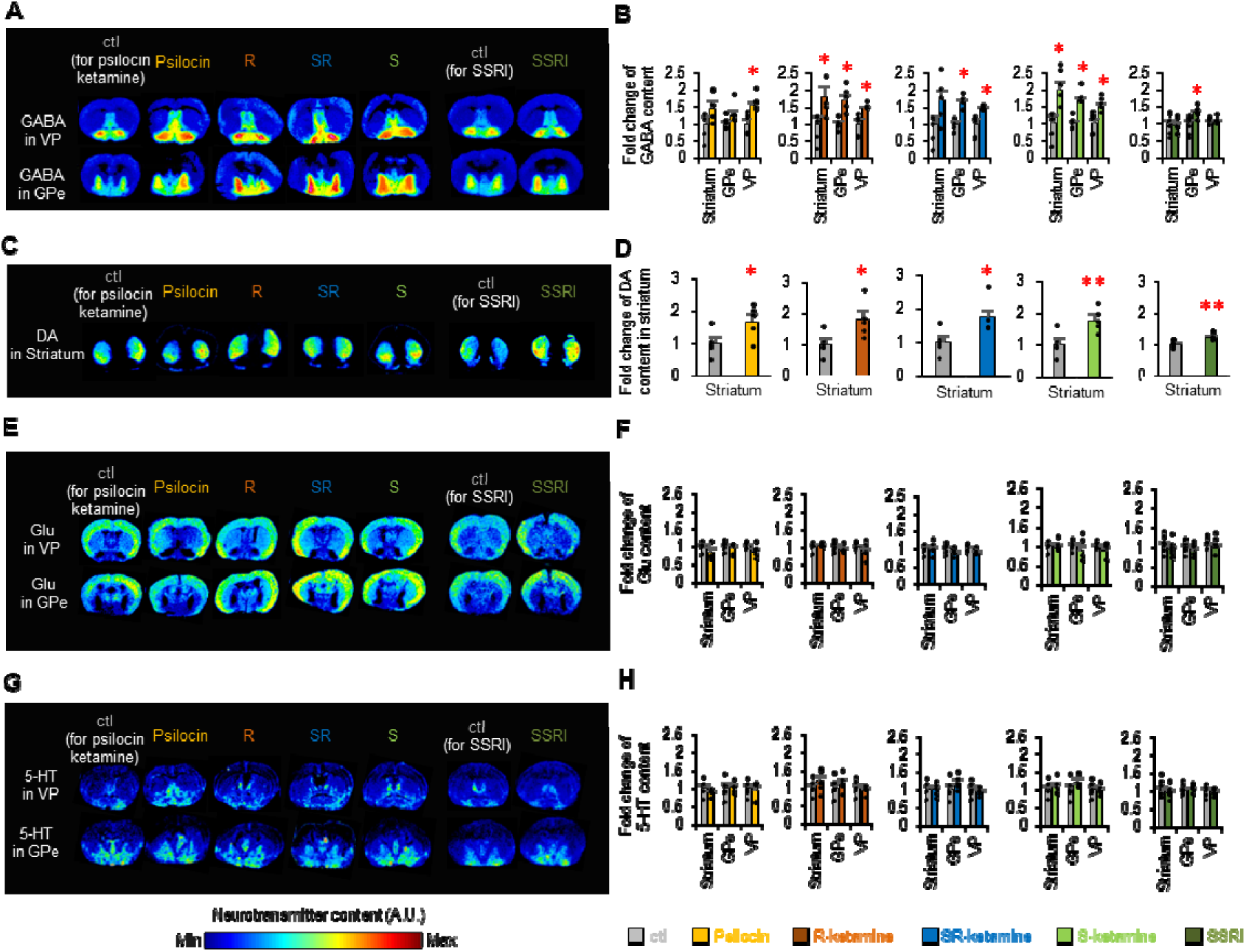
Pallidal γ-aminobutyric acid (GABA) and striatal dopamine (DA) are both increased across all antidepressant treatments. (A) Representative imaging mass spectrometry (IMS) images of GABA in the external globus pallidus (GPe) and ventral pallidum (VP) of treated and control (ctl) mice. (B) Fold changes in GABA content in the striatum, GPe, and VP are plotted (versus ctl mice). GABA content in the basal ganglia was increased after all of the indicated antidepressant treatments (n = 5 mice per treatment). (C, D) Representative IMS images of DA in the striatum of treated and ctl mice. DA content in the striatum was increased after all of the indicated antidepressant treatments (n = 5 mice per treatment). (E, F) Representative IMS images of glutamate (Glu) in treated and ctl mice. Glu content in the basal ganglia did not change after antidepressant treatments (n = 5 mice per treatment). (G, H) Representative IMS images of serotonin (5-HT) in treated and ctl mice. The 5-HT content in the basal ganglia did not change after antidepressant treatments (n = 5 mice per treatment). *p < 0.05. **p < 0.01 (Student’s t tests were used to compare treatments with ctls; p values are presented with post hoc Bonferroni corrections for multiple comparisons). The values are plotted as the means ± standard errors of the means.

Because IMS can provide a comprehensive analysis of neurotransmitters, we also examined the 5-HT, glutamate, and dopamine content from the same section. Surprisingly, the dopamine content in the striatum was upregulated after antidepressant treatment (Fig. 3C, D).

In addition, the glutamate content in the basal ganglia was not altered by any antidepressants (Fig. 3E, F). Although SSRIs and psilocin are 5-HT-related antidepressants, the 5-HT content in the basal ganglia was not altered by any antidepressants (Fig. 3G, H). These results indicate that antidepressant treatment increased dopamine (DA) levels and GABA levels in the basal ganglia.

### Antidepressants can be divided into two subtypes based on the patterns of volumetric change they induce

Using principal component analysis (PCA) and hierarchical clustering on the brain volume change patterns of the ECS and various antidepressant treatments compared with those of the control group, we found distinct differences between the antidepressant group and the ECS group (Fig. 4A, B). This result supported the distinctive features of antidepressant treatment and ECS (Fig. 1B–G). Despite the similarity among antidepressants, PCA of the volumetric changes at the selected time points (1 week post-injection for ketamine and psilocin; 3 weeks for SSRI) revealed two subgroups (Fig. 4A, B). The type I group comprised SSRI and S-ketamine, while the type II group included R-ketamine, SR-ketamine, and psilocin. This classification was supported further by the post hoc ROI-based analysis of the overlapping area assessment (Supplementary Fig. 3).

**Fig. 4:**
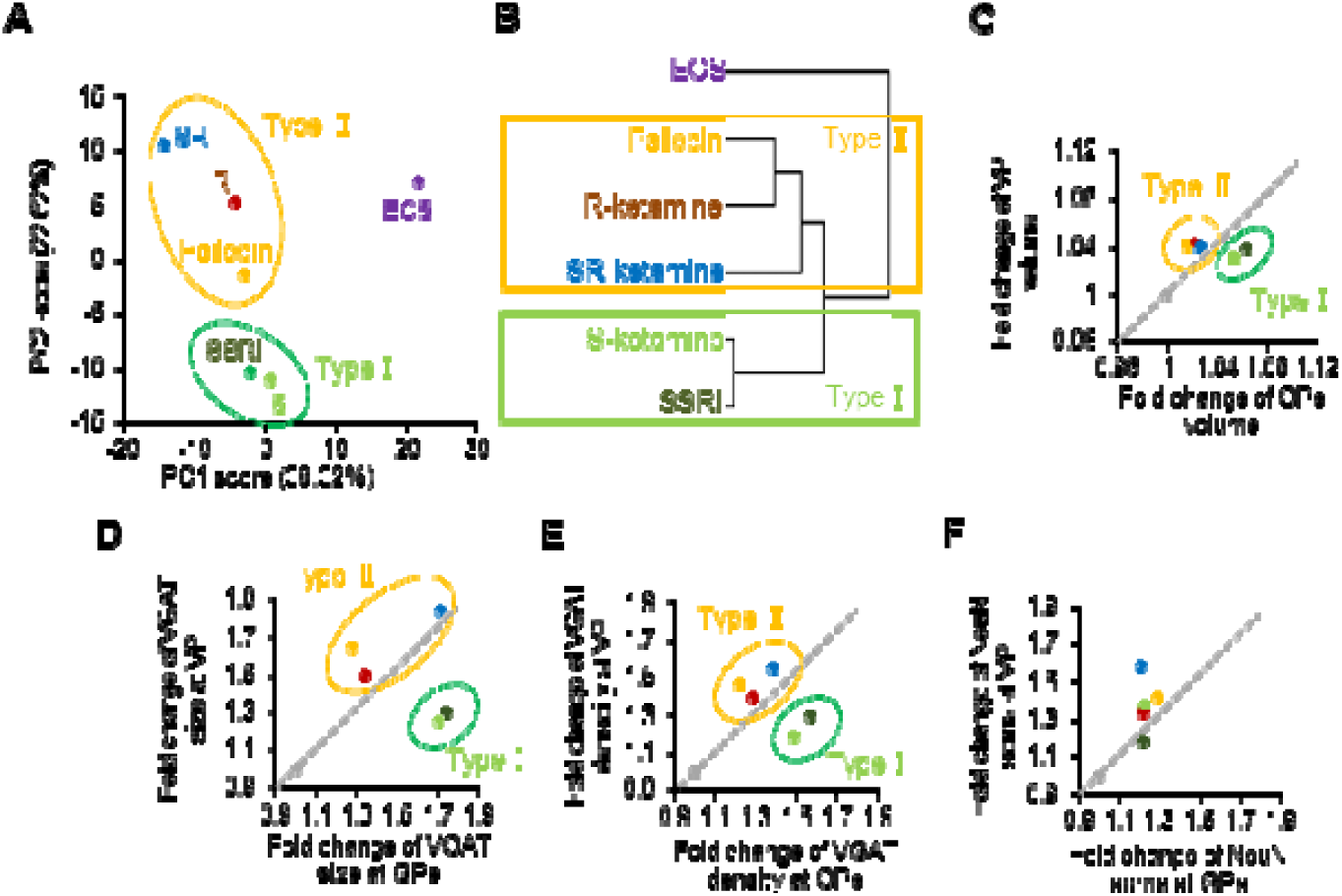
Principal component analysis identified two subgroups of volumetric changes. (A, B) Principal component analysis and hierarchical clustering of volume changes in treated mice. (C) The scatter plots of the fold change in the volume of the globus pallidus (GPe) and the ventral pallidum (VP) show VP predominant volume increases (type II group, yellow circles) and GPe predominant volume increases (type I group, green circles). (D, E) Scatter plots of the fold changes in the vesicular γ-aminobutyric acid (GABA) transporter (VGAT) puncta size and density and neuronal nuclei (NeuN) soma size in the GPe and VP. SR, *SR*-ketamine; R, *R*-ketamine; ECS, electroconvulsive stimulation; SSRI, escitalopram.

When examining the volumetric changes in the GPe and VP, we found that the enlargement indices (fold change relative to each control) of brain volume in the VP were greater than those in the GPe after treatment with type II medications (psilocin, *R*-ketamine, and *SR*-ketamine) and vice versa (Fig. 1I and Fig. 4C). This classification was consistent with the DBM results (Supplementary Fig. 4I). Interestingly, this macroscopic property was consistent with the microscopic property of MSN terminals but not soma size in both the VP and GPe principal neurons (Fig. 2C–E and Fig. 4D–F). This observation indicates that the type II group potentiates ventral MSN terminal hypertrophy, while the type I group potentiates dorsal MSN terminal hypertrophy.

### Volumetric structural covariance analysis highlights the NAc–VP–MD/LHb pathway

The VP mainly receives inhibitory inputs from the NAc and projects to the MD and LHb ^33–35^. In particular, the VP–LHb pathway has been implicated in depression ^34, 36^ and its antidepressant effect ^34, 37^. The GPe also receives inhibitory input mainly from the dorsal striatum and projects to the GPi and SNr. Our VBM analysis suggested that the volume increases in the VP and GPe act as “hub regions” in the context of antidepressant responses. To investigate this possibility, we conducted a structure covariance network analysis ^22, 23^, which allowed an unbiased examination of the correlation between volumetric changes in paired regions (Supplementary Fig. 7A). From these correlation matrices and network analyses, we found that the VP and MD were common hub regions in response to all classes of antidepressants (Supplementary Fig. 7B). The GPe was not identified as a hub region.

Next, to assess the relevance of volumetric changes in the VP, we conducted a target ROI-based structure covariance analysis between the VP and known connecting regions, including the dorsal striatum, caudate putamen (CPu), NAc, central amygdala (CeA), basolateral amygdala (BLA), GPe, MD, LHb, lateral hypothalamic area (LHA), lateral preoptic area (LPO), substantia nigra pars reticulata (SNr), ventral tegmental area (VTA), and midbrain reticular nucleus (MRN) ^34, 38, 39^. According to this VP-centered analysis, the difference in structural covariances (obtained by subtracting the Pearson correlation coefficient of the control from that of each antidepressant-treated group) were greatest for the VP–MD across all antidepressants (Supplementary Fig. 7C, E). By contrast, the differences were more modest for the NAc–VP and the VP–LHb (Supplementary Fig. 7D, F, G). These results suggest that antidepressant treatment strengthens structural connectivity within the NAc–VP–MD or NAc–VP–LHb pathways.

Next, we analyzed structural changes in the MD and LHb to support the observed volumetric reductions, focusing on excitatory type I/II vesicular glutamate transporter (VGluT1/2)^+^ and inhibitory VGAT^+^ presynaptic structures (Supplemental Fig. 8A, B). We found that across all antidepressants, the density but not the size of the VGAT^+^ puncta, decreased in the MD and LHb (Supplemental Fig. 8C, D). Additionally, in the MD, the density but not the size of VGluT1^+^ puncta, decreased with psilocin and *R*-ketamine treatment (Supplemental Fig. 8E, F). No structural changes were observed in the VGluT2^+^ puncta after treatment with any antidepressant (Supplemental Fig. 8G, H). These results indicate a common reduction in inhibitory terminals in the MD and LHb across all antidepressants, supporting the volumetric reductions observed in these regions.

### Striatal VGAT overexpression reproduces the structural and molecular changes associated with antidepressant-like effects

To address whether striatal VGAT overexpression reproduces the structural and molecular changes associated with antidepressant-like effects, we performed both gain- and loss-of-function experiments targeting VGAT in the NAc of wild-type mice. Specifically, we injected AAV5-hSyn-ALFA-VGAT and AAV5-CaMKII-EGFP-miR30-VGAT shRNA to overexpress and knock down VGAT, respectively, and used AAV5-hSyn-EGFP as a control. Four weeks after the AAV injection, we conducted behavioral and histological analyses (Fig. 5A). VGAT overexpression significantly reduced anxiety-like behaviors, whereas VGAT knockdown did not produce such effects (Fig. 5B–D).

**Fig. 5:**
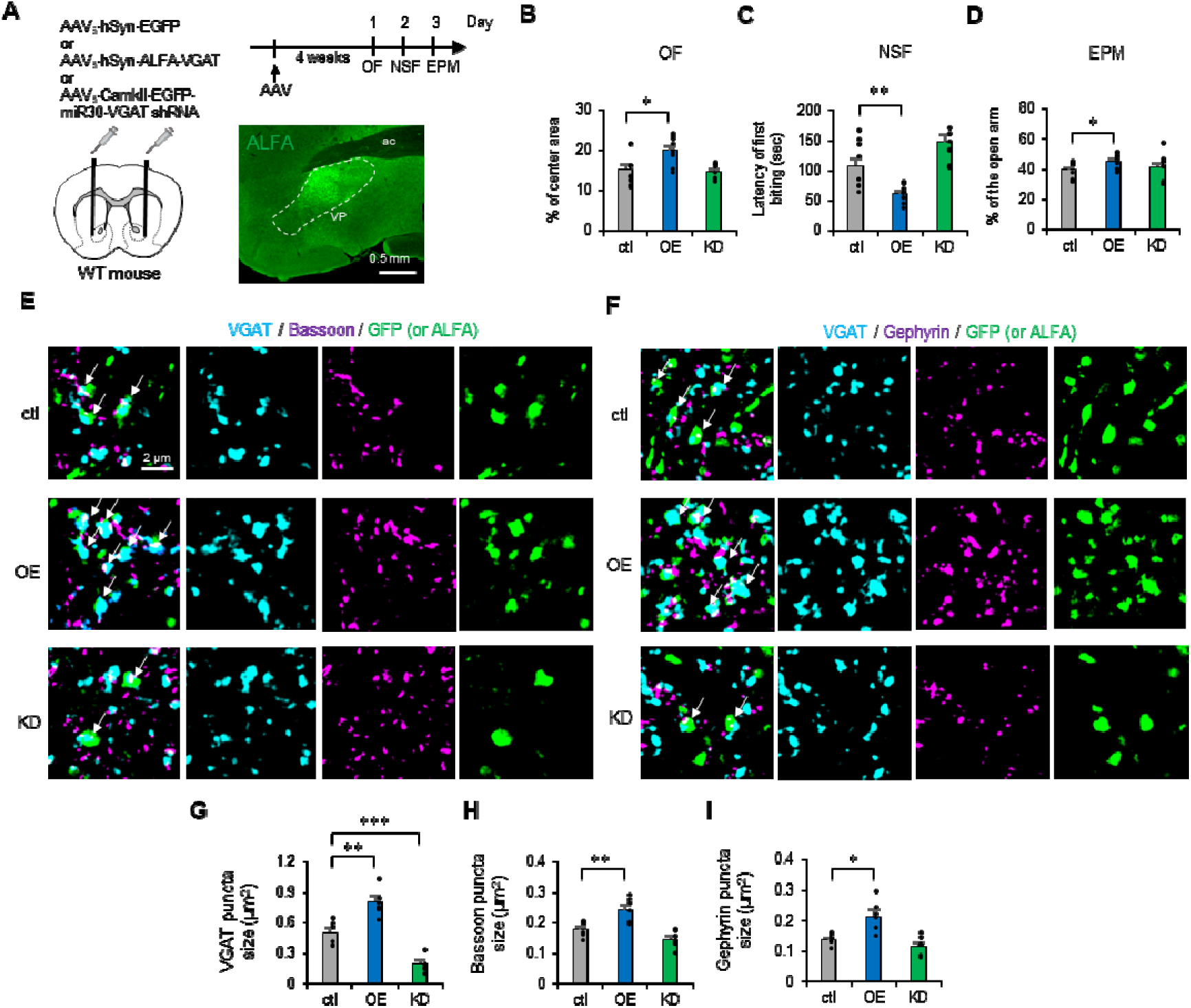
Overexpression of vesicular γ-aminobutyric acid transporter (VGAT) in nucleus accumbens (NAc) medium spiny neurons (MSNs) decreases innate anxiety and increases MSN terminal puncta. (A) The adeno-associated virus (AAV) vectors—AAV_5_-hSyn-EGFP for ctl, AAV_5_-hSyn-ALFA-VGAT for VGAT overexpression (OE), and AAV_5_-CamklJ-EGFP-miR30-VGAT shRNA for VGAT knockdown (KD)—were injected into the bilateral NAc of wild-type (WT) mice. A representative image shows ALFA staining in the ventral pallidum (VP) of OE mice. ac, anterior commissure. (B–D) Behavioral outcomes of the open field (OF), novelty suppressed feeding (NSF), and elevated plus maze (EPM) tests in ctl (n = 9), VGAT OE (n = 9), and VGAT KD (n = 9) mice. (E) Representative super-resolution microscopy images of VGAT, Bassoon, and green fluorescent protein (GFP; or ALFA) staining in the VP of ctl, VGAT OE, and VGAT KD mice. Arrows indicate GFP^+^ (or ALFA^+^) MSN terminals. (F) Representative super-resolution microscopy images of VGAT, Gephyrin, and GFP (or ALFA) staining in the VP of ctl, VGAT OE, and VGAT KD mice. (G–I) The sizes of GFP^+^ (or ALFA^+^) VGAT^+^, Bassoon^+^, and Gephyrin^+^ puncta in the VP were compared between the VGAT OE (n = 6) or VGAT KD (n = 6) mice and ctl mice (n = 6). *p < 0.05, **p < 0.01, ***p < 0.001 (Student’s t tests were used; p values were adjusted with post hoc Bonferroni corrections for multiple comparisons). The values are plotted as the means ± standard errors of the means.

Using SRM in the VP, we examined the size of puncta positive for VGAT and synaptic markers. VGAT overexpression increased the size of puncta positive for VGAT, the presynaptic marker Bassoon, and the postsynaptic marker Gephyrin (Fig. 5E, G–I). By contrast, although VGAT knockdown significantly decreased the size of VGAT^+^ puncta, it did not affect the size of Bassoon^+^ or Gephyrin^+^ puncta (Fig. 5F, G–I). Based on our previous research ^6^, these increases in the size of VGAT^+^, Bassoon^+^, and Gephyrin^+^ puncta are indicative of MSN terminal hypertrophy.

IMS analysis of the GPe revealed that VGAT overexpression in dorsal striatal neurons was associated with increased GABA levels (Supplementary Fig. 9A–C), consistent with the observed increases in VGAT^+^ and Bassoon^+^ puncta size in the GPe (Supplementary Fig. 9D–F).

Taken together, these results indicate that VGAT overexpression in striatal neurons induces MSN terminal hypertrophy and increases GABA levels in the pallidum; these alterations are associated with antidepressant-like effects. These findings suggest that the common action of antidepressants may involve VGAT overexpression in MSNs, leading to increased GABA levels and terminal hypertrophy.

### Pallidal volume increases after SSRI administration in depression model mice

Given that we demonstrated the structural and molecular consequences of antidepressant administration in naïve mice (Figs. 1–3), we next attempted to evaluate whether the effects of antidepressants on brain volume are similar in a mouse model of depression. We used a well-validated combination of disease model and effective medication; namely, a CORT-induced depression model and SSRI treatment ^18, 40, 41^. In the CORT-induced model, mice were administered CORT via drinking water for 3 weeks to induce depressive-like behaviors, followed by 3 weeks of adjunctive SSRI treatment (Fig. 6A). Behavioral tests (the OF, NSF, and EPM) confirmed that CORT-treated mice exhibited increased anxiety-like behaviors, and that adjunctive SSRI treatment reduced anxiety-like behaviors (Fig. 6B–D) as previously described ^41^.

**Fig. 6:**
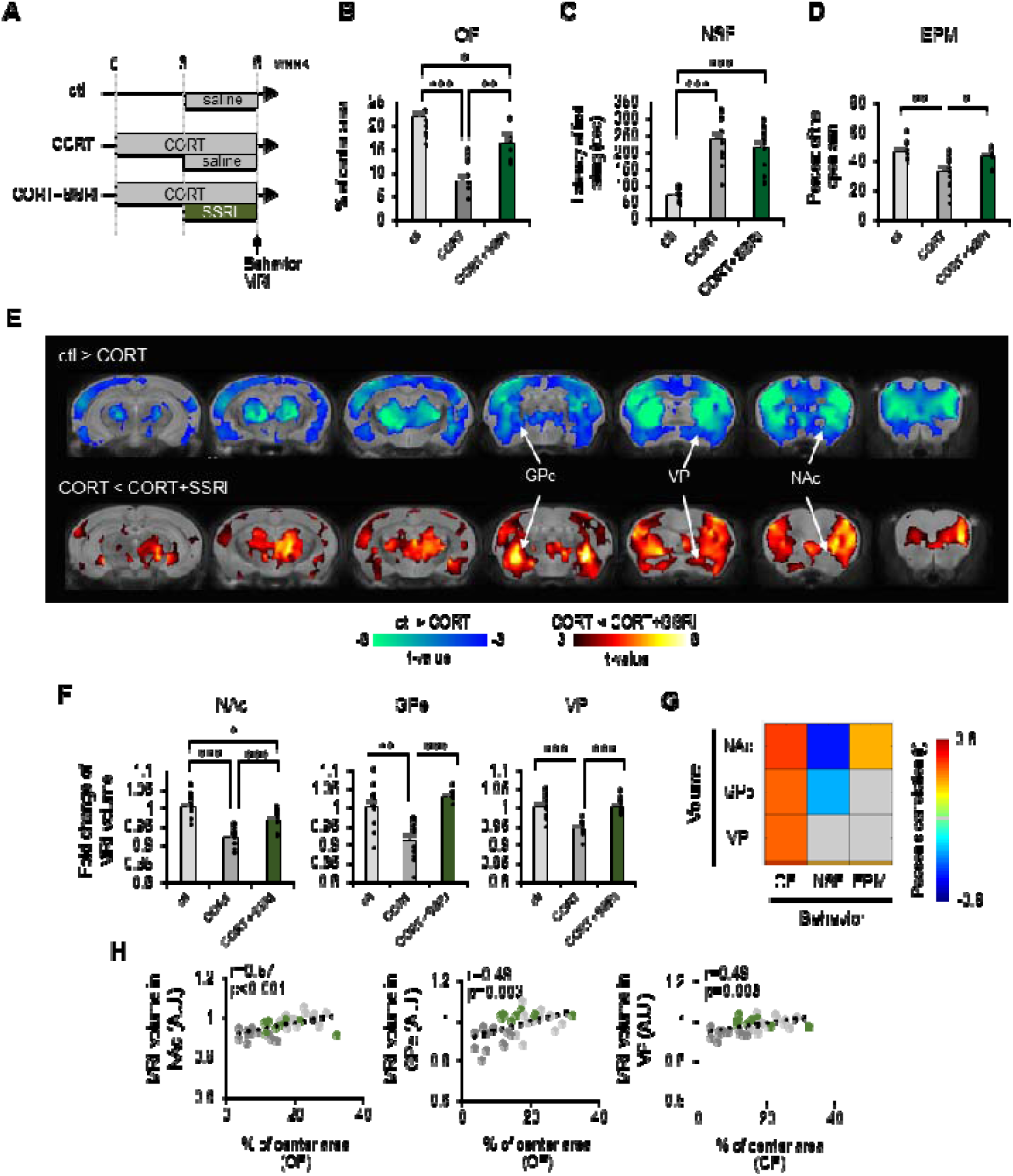
External globus pallidus (GPe) and ventral pallidum (VP) volumes also increase with selective serotonin reuptake inhibitor (SSRI) administration in corticosterone (CORT)-induced depression model mice. (A) Experimental time course for the administration of CORT and SSRI. (B–D) Behavioral outcomes of the open field (OF), novelty suppressed feeding (NSF), and elevated plus maze (EPM) tests in control (ctl) (n = 12), CORT-treated (n = 12), and CORT-treated mice with SSRI administration (n = 11). (E) Voxel-based morphometry (VBM) analysis comparing brain volume differences between ctl (n = 12) and CORT-treated mice (n = 12), and between CORT-treated mice and CORT-treated mice with SSRI (n = 11). Cool colors indicate significant volume decreases in CORT-treated mice compared with ctl mice (family-wise error corrected p < 0.05), and hot colors indicate significant volume increases in CORT-treated mice with SSRI compared with CORT mice. (F) Fold changes in the volumes of the nucleus accumbens (NAc), GPe, and VP. Colors indicate significant correlations between behavioral outcomes (OF, NSF, and EPM) and brain volumes (NAc, GPe, and VP). Gray color indicates no significant correlation. (H) Scatter plots showing OF outcome versus brain volumes (NAc, GPe, and VP). r refers to Pearson’s correlation coefficient. *p < 0.05, **p < 0.01, ***p < 0.001 (Student’s t tests were used; p values were adjusted with post hoc Bonferroni corrections for multiple comparisons). The values are plotted as the means ± standard errors of the means.

Subsequent VBM analysis revealed that in the CORT model, NAc, GPe, and VP volumes were reduced; however, SSRI treatment restored the volumes of these regions (Fig. 6E, F). These volume-increasing effects of SSRI in the depression model were similar to those in wild-type mice (Fig. 1B). Moreover, correlation analysis demonstrated that the volumetric changes in the NAc, GPe, and VP were significantly associated with behavioral outcomes (Fig. 6G). Collectively, these findings suggest that the degree of volume increase in the pallidum is linked to antidepressant-like effects in depression model mice.

## Discussion

This study demonstrated that the pallidal volume increase was a shared structural footprint of antidepressant treatment and that this increase was accountable for the enlargement of the MSN terminals with VGAT overexpression. Neurochemical analysis by IMS demonstrated an increase in GABA levels after antidepressant treatment. Artificial VGAT overexpression in MSNs was sufficient to reproduce the MSN terminal hypertrophy and an increase in GABA levels and was associated with antidepressant-like effects. Since we previously demonstrated that VGAT overexpression in MSNs enhanced GABA transmission at the pallidum ^6^, our findings suggest that a common mechanism of action of antidepressants involves enhanced GABA-mediated pallidal inhibition.

Structural covariance analysis revealed potential connectivity links between the VP and both its upstream and downstream structures. The VP receives input from the NAc and serves as an output nucleus of the basal ganglia, projecting to various regions involved in regulating motivated behavior, hedonic states, reinforcement, and reward/aversion processing ^33–35, 38^. The current data indicate that antidepressant treatments induce VGAT overexpression, increased GABA content, and presynaptic terminal hypertrophy in NAc MSNs. In our previous study, VGAT overexpression in dorsal striatal MSNs led to terminal hypertrophy, increased GABA content, and enhanced GABA transmission in the MSN output regions ^6^.

Moreover, we observed similar enhancements in GABAergic transmission within MSN output regions (such as the GPe and SNr) in L-DOPA-induced dyskinesia mouse models ^6^, which also exhibited MSN terminal hypertrophy, VGAT overexpression, and elevated GABA levels ^6^. On the basis of these converging lines of evidence, we propose that antidepressant-induced modifications in the NAc likely lead to enhanced GABAergic transmission to the VP, potentially resulting in VP inhibition.

Notably, although these upstream changes suggest an increased inhibitory drive to the VP, antidepressant treatments were consistently associated with selective decreases in MD and LHb volumes alongside an increase in VP volume (Supplementary Fig. 7E, F). The LHb volume reduction is particularly intriguing given that heightened LHb activity contributes to depressive states and both SSRIs and ketamine suppress LHb activity ^34, 36^. We need to acknowledge that the causal relationship between the suppression of LHb activity and the decrease in LHb volume is currently unknown, and we only identified that the antidepressant-induced decrease in LHb volume coincides with decreased inhibitory presynapse density but not with decreased excitatory presynaptic density (Supplementary Fig. 8). Such histological data imply the disinhibition of LHb activity after antidepressant treatment, which seems contradictory to the prevailing view ^34, 36^. To understand the functional and structural consequences of LHb more fully, at the very least, analyses of animal models of depression subjected to antidepressant treatments are required. In addition, the engagement of VP–LHb monosynaptic or polysynaptic connections should be clarified to determine the relevance of the NAc–VP–LHb pathway.

VP projections to the MD are important for the cognitive component of reward learning ^42^, and MD activity is associated with stress resilience in mice ^43^, indicating the role of MD in depression pathophysiology. Human imaging studies have demonstrated a decrease in MD volume in individuals with stress-related disorders, including depression ^44^ and posttraumatic stress disorder ^45^. However, other thalamic nuclei in addition to the MD showed a decreased volume in these disorders. The antidepressant effect on thalamic volume was specific to the MD in mice, suggesting a distinct mechanism for the decrease in MD volume after antidepressant treatment. The biological importance of the decreased MD volume and the NAc–VP–MD pathway could be addressed by the abovementioned approach as it was for the LHb.

Recent studies have provided compelling evidence that SSRIs, ketamine, and psilocybin share a common binding site within the brain-derived neurotrophic factor (BDNF) receptor, tropomyosin receptor kinase B (TrkB) ^9, 10^. Given that striatal MSNs are highly sensitive to BDNF, which induces the selective hypertrophy of MSN dendrites ^46^, it is conceivable that the activation of TrkB signaling by these diverse antidepressants converges on similar downstream mechanisms. Such TrkB-dependent pathways may potentiate neuronal plasticity and drive the observed structural remodeling of MSN presynaptic terminals. Despite their distinct primary pharmacological actions, the convergence on TrkB-mediated plasticity may underlie the parallel structural outcomes observed in the present study, thereby providing a unified framework for understanding the common antidepressant effects across these drugs.

Based on the patterns of volume increase in the GPe and VP, we found that antidepressants can be divided into two subgroups. This subdivision is not only reflected in brain volume changes but also coupled with structural changes in MSN terminals (Fig. 4). It should be noted that the subdivisions clearly separated the chronic volumetric effects by *R*- or *S*-ketamine. Previous studies supported this distinction; functional MRI studies with rats showed distinct acute activation patterns between *R*-ketamine and *S*-ketamine ^47^.

[^18^F]-Fluorodeoxyglucose positron emission tomography of human brains revealed a distinct acute metabolic pattern between *R*-ketamine and *S*-ketamine ^48^. c-Fos mapping with mouse brains also revealed distinct activation patterns between *R*-ketamine and *S*-ketamine ^49^.

Although the difference in the sustained effects is not clear in humans, *R*-ketamine showed more sustained antidepressant effects than *S*-ketamine in animals ^15^. Combined with the sustained antidepressant effect of psilocybin in humans ^5^, the grouping with *R*-ketamine and psilocin may be related to their long-term effects. In contrast, the grouping with SSRIs and *S*-ketamine may not be.

We consistently observed a delayed increase in dopamine content in the basal ganglia following antidepressant treatment (Fig. 3C, D). Previous studies have demonstrated that ketamine and psilocin acutely elevate extracellular dopamine concentrations in the medial prefrontal cortex, NAc, hippocampus, and amygdala via activation of serotonin and glutamate neurons ^50, 51^. However, our results did not show a chronic increase in serotonin or glutamate content in the basal ganglia. It is important to note that our IMS analysis measured total tissue dopamine without distinguishing between the intracellular and extracellular pools ^52^. Our findings thus suggest that further investigations into dopamine synthesis and metabolism—and not merely dopamine neuron activity—are needed to elucidate the mechanisms underlying the therapeutic actions of antidepressants. Furthermore, the relevance of increased dopamine content in the basal ganglia has remained unclear, but it is expected to clarify whether chronic increases in dopamine content are related to antidepressant effects.

In summary, we discovered shared structural and molecular changes after antidepressant treatments. An increase in pallidal volume may serve as a structural biomarker indicating the efficacy of antidepressants. The antidepressant-induced increase in GABA-mediated inhibition of the pallidum can be attributed to VP-targeted neuromodulation to mimic the therapeutic effect. An increased GABA content in the pallidum may serve as a therapeutic biomarker that can be detected using magnetic resonance spectroscopy.

## Acknowledgments

We thank Collaborative Research Resources, Keio University School of Medicine, for technical assistance with the Zeiss ELYRA 3D-SIM system. We thank Dr. Megumu Takahashi and Dr. Hiriyuki Hioki (Juntendo University Graduate School of Medicine, Tokyo, Japan) for helping to make the AAV. We also thank Lisa Kreiner, PhD, and Bronwen Gardner, PhD, from Edanz (https://jp.edanz.com/ac) for editing the English text of a draft of this manuscript.

## Author contributions

K.F.T. supervised the project; Y.A. and K.F.T. designed the experiments; Y.A. and S.Y. performed and analyzed the MRI studies; Y.A. performed and analyzed the histological assessments; Y.A., R.M., S.T., and Y.S. performed and analyzed the imaging mass spectroscopy; S.M. made AAV; K.Y. and D.I. provided psilocin; K.H. provided ketamine; and Y.A. and K.F.T. wrote and revised the original manuscript.

## Data availability

All data are available in the main text or the supplementary materials. Materials, resources, plasmids, computer codes, and reagents are available from the corresponding author upon request.

