## Supplemental information for "Distinct classes of antidepressants commonly act to shape pallidal structure and function in mice"

**Supplementary Figures**

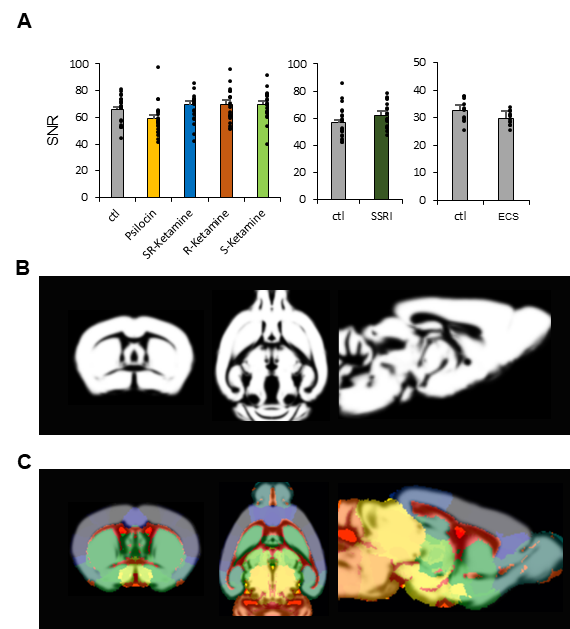

**Supplementary Fig. 1: Quality check of magnetic resonance imaging (MRI) data and analysis.**

(A) Signal-to-noise ratios (SNRs) of all MRI images were plotted. (B) Representative results of the gray matter after SPM segmentation. (C) Overlaid images of the Advanced Normalization Tools (ANTs) registered brain atlas and the gray matter images. Ctl, control; SSRI, selective serotonin reuptake inhibitor; ECS, electroconvulsive stimulation.

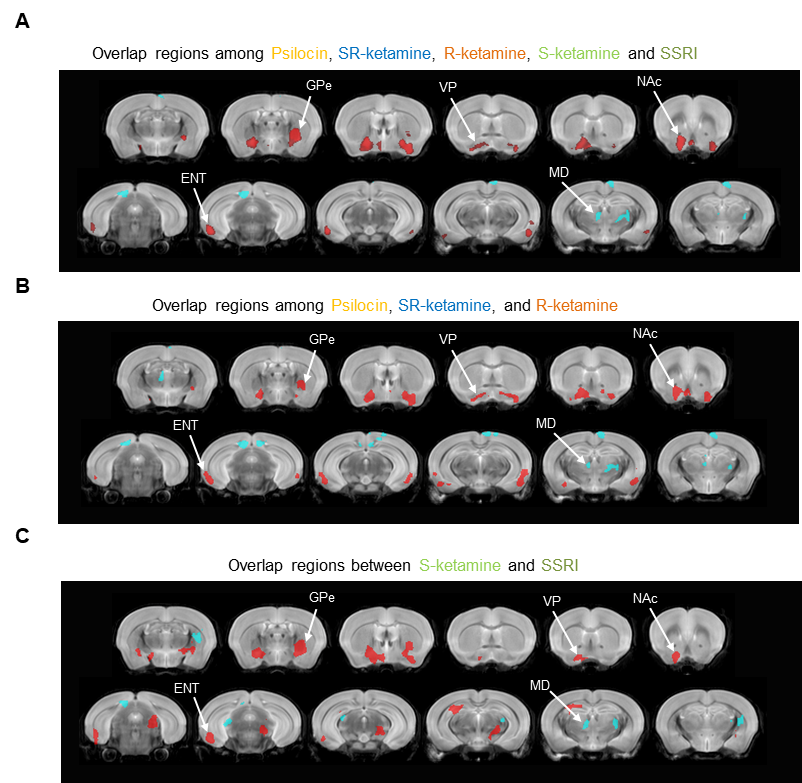

**Supplementary Fig. 2: Overlapping regions of brain volume changes in voxel-based analysis.**

(A) Overlapping regions of brain volume changes among all treated mice. This result is the same as that in Figure 1H. (B) Overlapping regions among psilocin, *SR*-ketamine, and *R*-ketamine. (C) Overlapping regions between *S*-ketamine and escitalopram (SSRI). Red indicates commonly increased volumes, and blue indicates commonly decreased volumes. ENT, entorhinal cortex; GPe, globus pallidus; MD, mediodorsal nucleus of the thalamus; NAc, nucleus accumbens; VP, ventral pallidum.

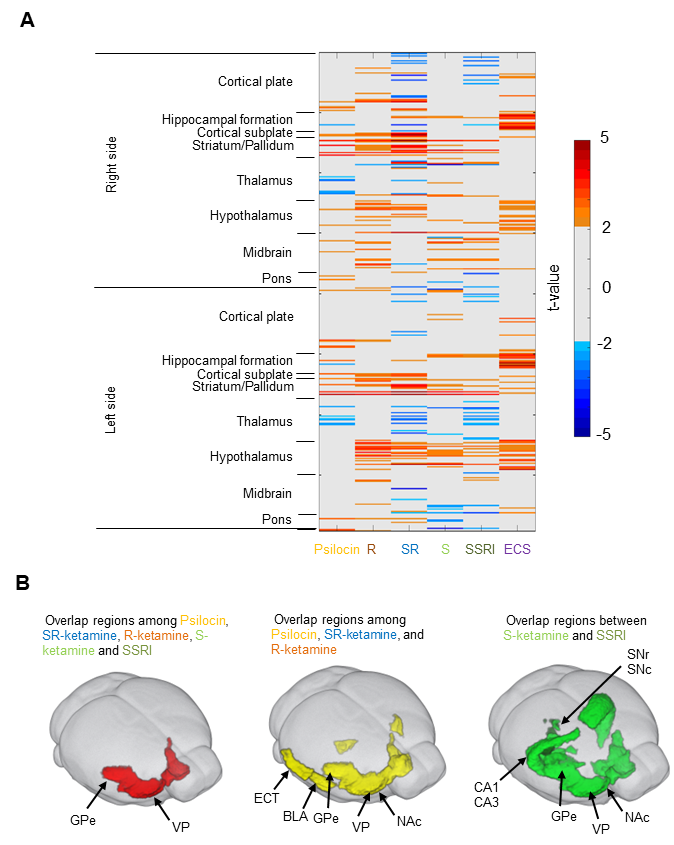

**Supplementary Fig. 3: ROI-based analysis in each type of treated mouse.**

(A) Region of interest (ROI)-based analysis of brain volume differences between treated mice and appropriate control mice (ctl). Hot colors indicate significant increases in volume compared with those in the ctl mice (uncorrected *p* < 0.01), and cool colors indicate the opposite. (B) Overlapping regions in the ROI-based analysis results. Red indicates overlapping regions among all treated mice. Yellow indicates overlapping regions among psilocin, *SR*-ketamine, and *R*-ketamine-treated mice. Green indicates overlapping regions between mice treated with *S*-ketamine and those treated with escitalopram (SSRI). BLA, basolateral amygdala; CA1, hippocampal cornu ammonis 1 (CA1) region; CA3, hippocampal CA3 region; ECT, electroconvulsive therapy; ENT, entorhinal cortex; GPe, globus pallidus; NAc, nucleus accumbens; SNc, substantia nigra pars compacta; SNr, substantia nigra pars reticulata; VP, ventral pallidum.

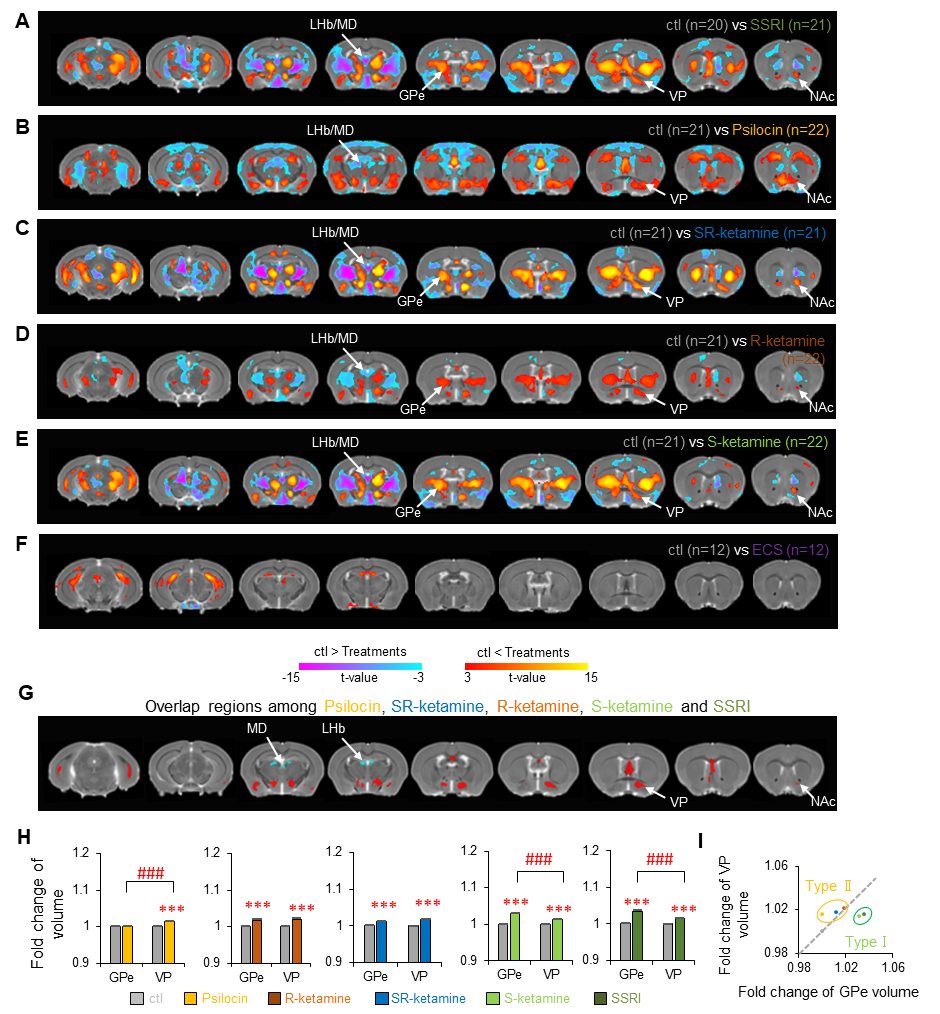

**Supplementary Fig. 4: Voxel-based morphometry (VBM) results were validated using template-free registration deformation-based morphometry (DBM) analysis.**

(A–F) DBM was performed to compare brain volume differences between treated mice and appropriate control (ctl) mice. Hot colors indicate significant volume increases compared with the ctl mice (family-wise error corrected p < 0.05) and cool colors indicate the opposite (i.e., significant volume decreases). (G) Volume changes in overlapping regions of the brain after treatment. Red indicates commonly increased volumes and blue indicates commonly decreased volumes. (H) Fold changes in the volumes of the external globus pallidus (GPe) and ventral pallidum (VP) after each antidepressant treatment. ***p < 0.001 (Student’s t tests were used for comparisons with ctl mice; p values were adjusted with post hoc Bonferroni corrections for multiple comparisons). ###p < 0.001 (Student’s t tests were used for comparisons between the GPe and VP of treated mice, p values are presented with post hoc Bonferroni corrections). (I) Scatter plots of the fold changes in the volumes of the GPe and VP reveal VP-predominant volume increases (type II group, yellow circles) and GPe-predominant volume increases (type I group, green circles). LHb, lateral habenula; MD, mediodorsal nucleus of the thalamus; NAc, nucleus accumbens; SSRI, selective serotonin reuptake inhibitor.

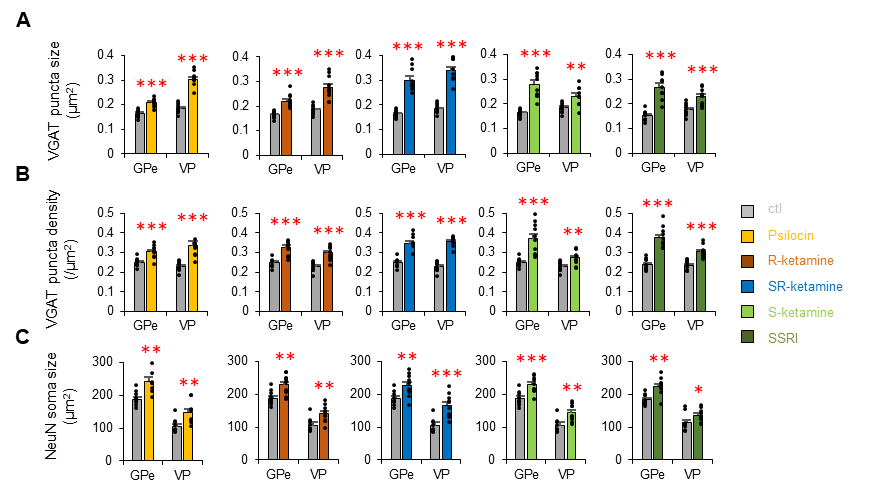

**Supplementary Fig. 5: The structural pattern changes of MSN terminals.**

(A–C) The size and density of vesicular γ-aminobutyric acid (GABA) transporter (VGAT) puncta of medium spiny neuron (MSN) terminals and the soma size of globus pallidus (GPe)/ventral pallidum (VP) neurons relative to each control (ctl) were compared in the GPe and VP of antidepressant-treated mice (n = 10 mice for each treatment). **p* < 0.05, ***p* < 0.01, ****p* < 0.001 (Student *t* tests was used for comparisons between the GPe and VP). The values are plotted as the means ± standard errors of the means. SSRI, escitalopram.

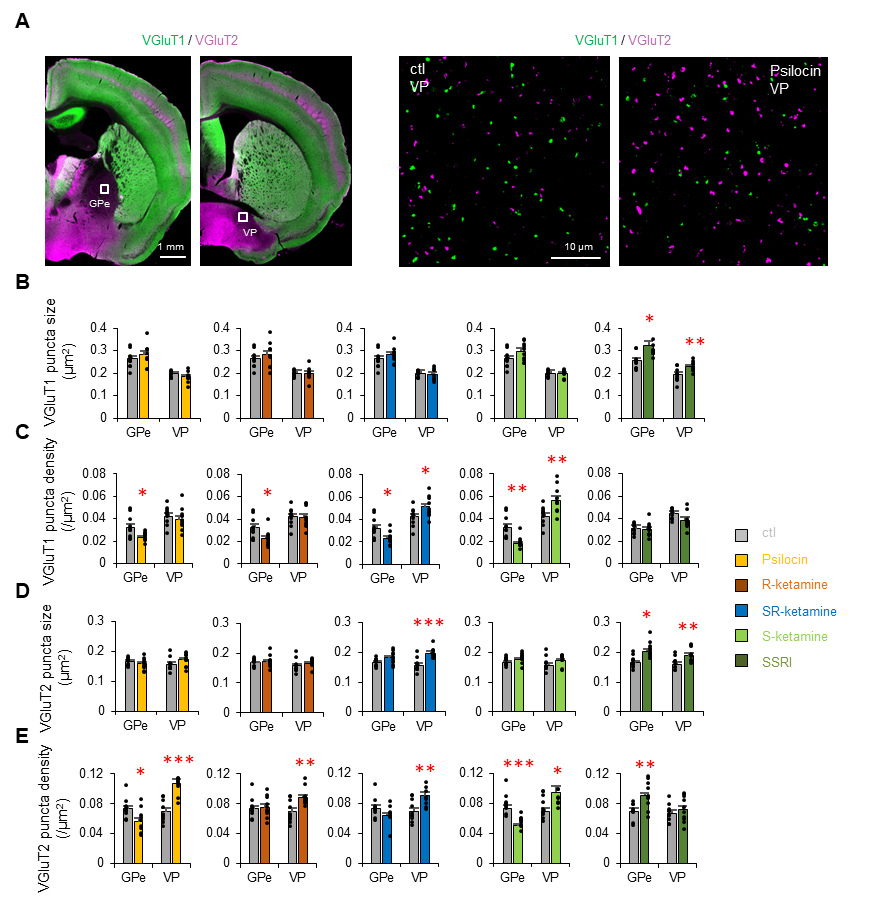

**Supplementary Fig. 6: Structural changes in excitatory terminals in the GPe and VP of antidepressant-treated mice.**

(A) Representative microscopy images of type I vesicular glutamate transporter (VGluT1), and type II VGlu (VGluT2) staining in the globus pallidus (GPe) and the ventral pallidum (VP). Representative super-resolution microscopy images of VGluT1 and VGluT2 staining in the VP of control (ctl) and psilocin-treated mice are shown. VGluT1 is an excitatory terminal marker of cortical neurons, and VGluT2 is an excitatory terminal marker of thalamic neurons. (B–E) The size and density of VGluT1 and VGluT2 puncta in the GPe and VP were compared between treated and ctl mice (n = 10 mice for each treatment). **p* < 0.05, ***p* < 0.01, ****p* < 0.001 (Student *t* tests were used for comparisons, *p* values are presented with post hoc Bonferroni corrections for multiple comparisons). The values are plotted as the means ± standard errors of the means.

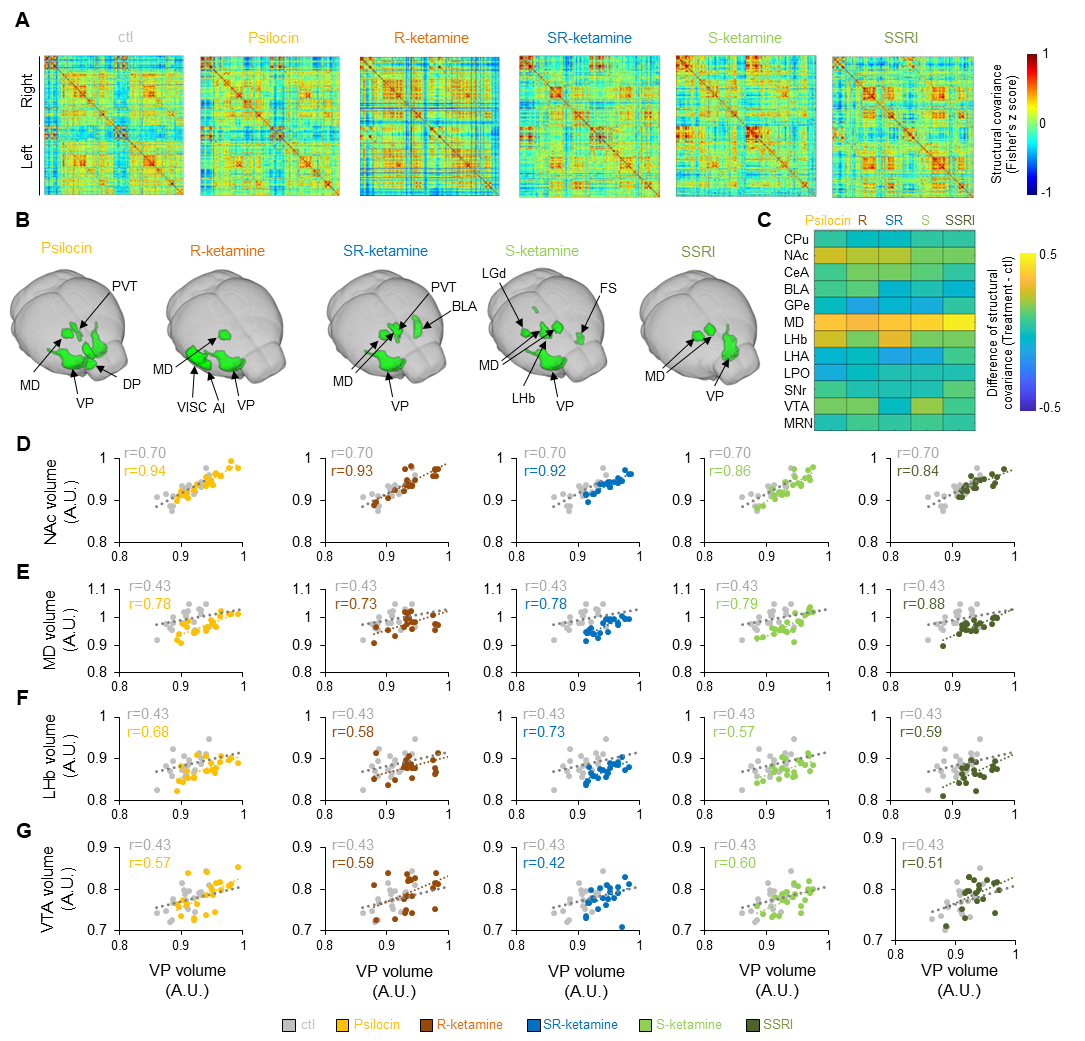

**Supplementary Fig. 7: Structural covariance analysis highlights the NAc–VP–MD/LHb pathway.**

(A) Region of interest (ROI)-based structural covariance matrices in antidepressant-treated mice. (B) Hub regions were calculated from structural covariance differences between treated and control (ctl) mice. The ventral pallidum (VP) and mediodorsal thalamus (MD) were common hub regions in all treated mice. (C) Differences in the structural covariance between the VP and connecting regions compared with those in ctl mice. (D–G) Plots of each brain volume and the VP volume in antidepressant-treated mice. Structural covariances of each region are shown as Pearson correlations (*r*). A.U., arbitrary units; AI, agranular insular cortex; BLA, basolateral amygdala; CeA, central amygdala; CPu, caudate putamen; DP, dorsal peduncular area; ENT, entorhinal cortex; FS, fundus of the striatum; GPe, globus pallidus, Hip, hippocampus; LGd, dorsal part of the lateral geniculate complex; LHA, lateral hypothalamus area; LHb, lateral habenula; LPO, lateral preoptic area; MRN, midbrain reticular nucleus; NAc, nucleus accumbens; PVT, paraventricular nucleus of the thalamus; SNr, substantia nigra pars reticulata; SSRI, escitalopram; VISC, visual cortex; VTA, ventral tegmental nucleus.

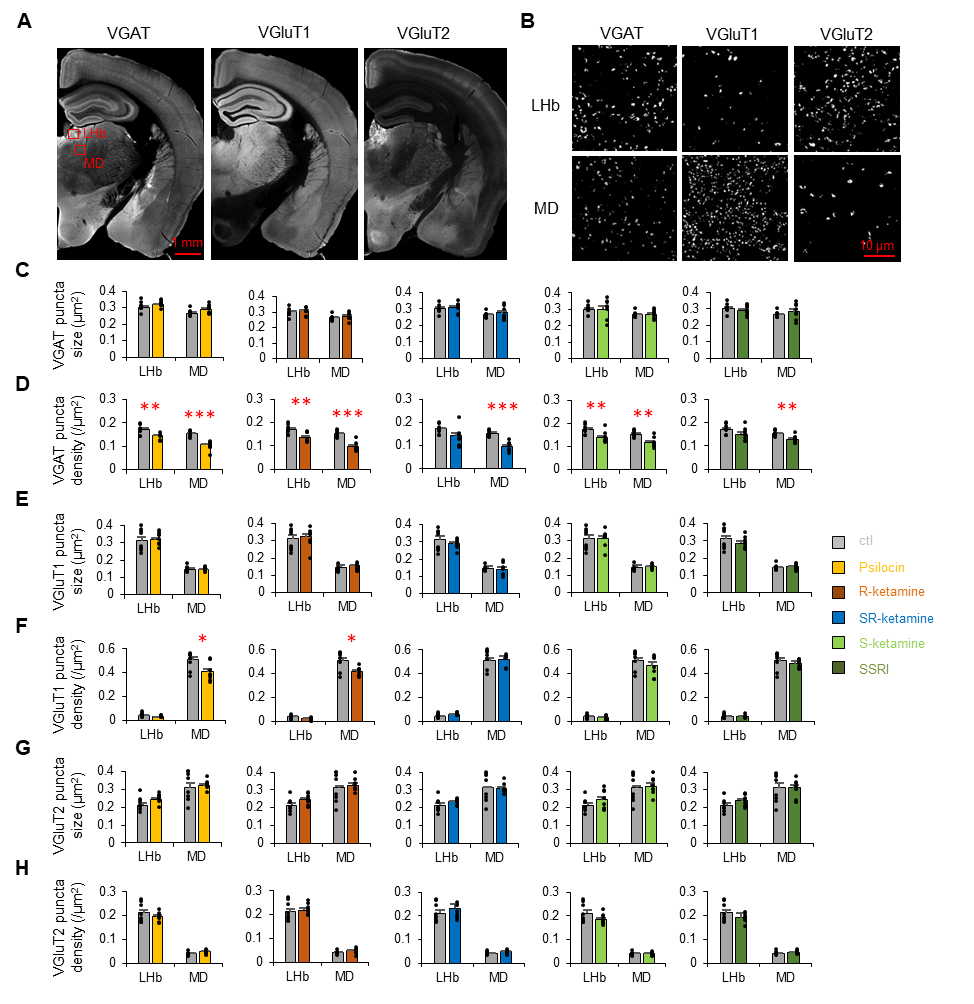

**Supplementary Fig. 8: Structural changes in inhibitory terminals in the LHb and MD of antidepressant-treated mice.**

(A) Representative microscopy images of vesicular γ-aminobutyric acid (GABA) transporter (VGAT), type I vesicular glutamate transporter (VGluT1), and type II VGlu (VGluT2) staining. The red squares of the lateral habenula (LHb) and mediodorsal thalamus (MD) are shown after further super-resolution microscopy (SRM) analysis. (B) Representative SRM images of VGAT, VGluT1, and VGluT2 staining in the MD and LHb. (C–H) The size or density of VGAT, VGluT1, or VGluT2 puncta in the LHb and MD were compared between treated and control (ctl) mice (n = 8 mice for each treatment). **p* < 0.05, ***p* < 0.01, ****p* < 0.001 (Student *t* tests were used to compare treatment and ctls, *p* values are presented with post hoc Bonferroni corrections for multiple comparisons). The values are plotted as the means ± standard errors of the means. SSRI, escitalopram.

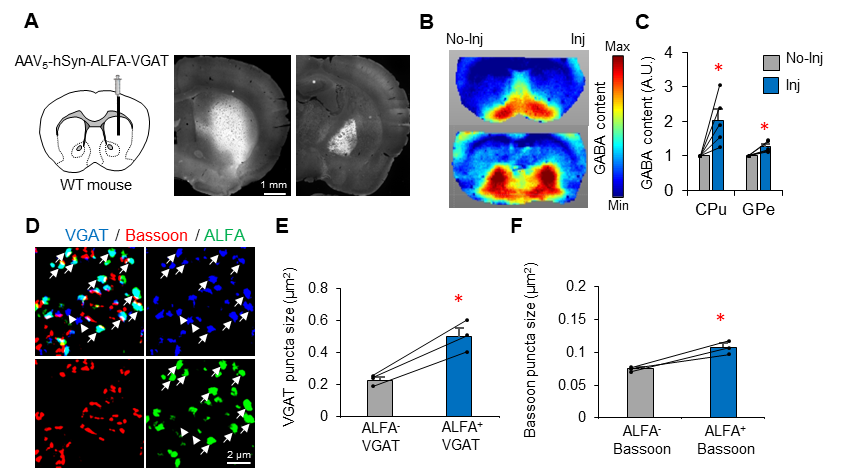

**Supplementary Fig. 9: Striatal overexpression of VGAT induced a shared biochemical responses of increased pallidal GABA content.**

(A) The AAV vectors of AAV_5_-hSyn-ALFA-VGAT-pA was injected into the dorsal striatum of wild-type (WT) mice. Representative image of ALFA staining. (B) Representative imaging mass spectrometry of GABA content in VGAT-overexpressing mice. (C) GABA content in the caudate putamen (CPu) and GPe were compared between the AAV-injected side (Inj) and the noninjected side (No-Inj) (n = 5). (D) Representative super-resolution microscopy images of VGAT, Bassoon (presynaptic protein), and ALFA in the GPe. The arrows indicate VGAT overexpressing terminals, and the arrowheads indicate non-overexpressing terminals. (E, F) The sizes of VGAT and Bassoon puncta in medium spiny neuron (MSN) terminals were compared between ALFA^+^ and ALFA^−^ puncta in the GPe (n = 3). *p < 0.05 (Student t tests were used with p values adjusted with post hoc Bonferroni corrections for multiple comparisons). The values are plotted as the means ± standard errors of the means.
